# The early evolution of oral poliovirus vaccine is shaped by strong positive selection and tight transmission bottlenecks

**DOI:** 10.1101/2020.08.20.260075

**Authors:** Andrew L. Valesano, Mami Taniuchi, William J. Fitzsimmons, Md Ohedul Islam, Tahmina Ahmed, Khalequ Zaman, Rashidul Haque, Wesley Wong, Michael Famulare, Adam S. Lauring

## Abstract

The evolution of circulating vaccine-derived polioviruses (cVDPV) from components of the live-attenuated oral poliovirus vaccine (OPV) presents a major challenge to global polio eradication. This process has largely been characterized by consensus sequencing of isolates collected from routine surveillance, and little is known about the early evolution of OPV within vaccinated hosts. These early events are critical steps in the progression of OPV to cVDPV. Here, we use whole genome, high depth of coverage sequencing to define the evolutionary trajectories of monovalent type 2 OPV in a cluster-randomized trial of polio vaccines in Matlab, Bangladesh. By sequencing 416 longitudinal samples from 219 mOPV2 recipients and 81 samples from 52 household contacts, we were able to examine the extent of convergent evolution in vaccine recipients and track the amount of viral diversity transmitted to new hosts. Using time-series data from a synchronized point of vaccine administration, we identify strong positive selection of reversion mutations at three known attenuating sites within two months post-vaccination. Beyond these three recognized “gate-keeper” mutations, we identify 19 mutations that exhibit significant parallelism across vaccine recipients, providing evidence for early positive selection not previously detected by phylogenetic inference. An analysis of shared genetic variants in samples from vaccinated individuals and their household contacts suggests a tight effective bottleneck during transmission. The absence of positively selected variants among household contacts across the cohort suggests that this tight bottleneck limits the transmission of these early adaptive mutations. Together, our results highlight the distinct evolutionary dynamics of live attenuated virus vaccines and have important implications for the success of novel OPV2 and other next generation approaches.

**Significance:** The emergence of circulating vaccine-derived polioviruses (cVDPV) through evolution of the oral polio vaccine (OPV) poses a significant obstacle to global eradication. Understanding the genetic changes in OPV that occur as it evolves and transmits in populations is important for preventing future cVDPV outbreaks. Little is known about the early events in VDPV evolution and the selective forces that drive them. We used high depth-of-coverage genome sequencing to assess the within-host evolutionary dynamics of monovalent type 2 OPV in a vaccine trial in Matlab, Bangladesh. We leverage longitudinal sampling from vaccine recipients and household contacts to identify mutations that arise in parallel across individuals and estimate the size of the transmission bottleneck. We find evidence for strong positive selection on key sites in the capsid and the 5’ noncoding region, many of which have not been previously identified. Our results also suggest that narrow transmission bottlenecks can constrain the spread of mutations selected within individuals. These results provide important insights into how OPV variants spread in populations and are highly relevant for ongoing poliovirus surveillance and the design of improved polio vaccines.

## Introduction

Genetic reversion and the associated loss of attenuation in oral poliovirus vaccine (OPV) strains are major barriers to achieving global poliovirus eradication (1). In areas of low vaccine coverage, OPV can evolve into circulating vaccine-derived polioviruses (cVDPV) that cause cases of poliomyelitis that are indistinguishable from those caused by wild polioviruses (WPV) (2–4). Of the three OPV serotypes, the Sabin type 2 is responsible for most cVDPV outbreaks to date (5, 6). Following the eradication of wild type 2 polioviruses, the Global Polio Eradication Initiative switched routine immunization schedules from trivalent OPV (tOPV, Sabin types 1, 2, and 3) to bivalent OPV (Sabin types 1 and 3) to reduce the risk of future cVDPV2 outbreaks (7). However, monovalent type 2 OPV (mOPV2) is still used to combat active cVDPV2 outbreaks. While the global replacement of tOPV with bOPV has reduced the presence of OPV2 in surveillance samples, cVDPV2 outbreaks remain a major problem. This is partly due to the continued reliance on monovalent Sabin type 2 OPV (mOPV2) to control new cVDPV2 outbreaks. Nearly half of the cVDPV2 outbreaks observed after the withdrawal of tOPV resulted from a previous mOPV2 intervention response (8).

The emergence of cVDPV is a recurrent evolutionary process that exhibits a high degree of parallel, or convergent, evolution. Most data on cVDPV come from poliovirus isolates in cases of acute flaccid paralysis or environmental surveillance (9–12). A recent study of type 2 cVDPV sequences from multiple outbreaks in five countries identified a limited number of sites under positive selection across independent lineages (9). Three mutations – A481G, U2909C (VP1-I143T), and U398C – were inferred to be under the strongest selection pressure and precede subsequent substitutions. The A481G and U398C mutations are located in the 5’ noncoding region and are functionally important to RNA structures in the internal ribosome entry site (IRES). All three mutations are known molecular determinants of attenuation, occur within the first two months after vaccination, and are associated with increased virulence in animal models (9, 10, 13–16). For these reasons, they are referred to as “gatekeeper” mutations that initiate the process of attenuation loss. Although phylogenetic studies have provided important information on the evolutionary trajectories of cVDPV, they are limited in their ability to resolve the exact timing of gatekeeper mutations and may lack power to detect natural selection due to sampling bias (17). Isolates of cVDPV have undergone months or years of evolution prior to isolation and lack a definitive link to the time of vaccine administration, further limiting our understanding of the early evolution of OPV in humans.

Investigating the evolutionary dynamics within individual hosts can complement phylogenetic studies of virus evolution (18). Complex evolutionary processes take place within the span of a single infection that cannot be resolved by standard consensus sequencing. Individual mutations, most often single nucleotide variants, arise within infected hosts and change in frequency according to the forces of natural selection and genetic drift. Studying how viruses evolve at this scale can uncover genomic sites under selective pressure, clarify the relative roles of selection and drift in viral evolution, and can inform sequence-based diagnostic and surveillance tools (19, 20).

Various sequencing approaches have been used to study the molecular epidemiology of poliovirus and to monitor OPV stocks for reversion (21, 22), but few have been purposed for measuring viral diversity within naturally infected hosts. Routine surveillance for VDPV involves sequencing only the region encoding the capsid protein VP1 (23). Many approaches for whole genome sequencing rely on amplification of viral isolates in cell culture, which may not accurately preserve the diversity present in the original specimen (24). Other high-throughput sequencing approaches that aim to measure within-host diversity have targeted only a specific portion of the poliovirus genome (25). While some have sequenced viral genomes or specific genomic regions from asymptomatic vaccine recipients (9, 26–28), we lack a comprehensive characterization of the early evolutionary dynamics of OPV within vaccinated individuals and during transmission to their close contacts.

Here we use whole genome, deep sequencing of stool samples from a clinical trial of OPV to elucidate the early evolution of polioviruses within and between human hosts. We developed an approach for sequencing OPV genomes directly from primary stool samples and validated its accuracy for identification of intrahost single nucleotide variants (iSNV). We applied this approach to samples from a recent trial that investigated the effect of tOPV cessation on the transmission of type 2 OPV (29). The trial included a defined point of introduction of monovalent type 2 OPV (mOPV2) and weekly longitudinal sampling of vaccine recipients and their household contacts; it therefore represents an opportunity to investigate the early evolutionary dynamics of OPV2 in a community setting. We identify several mutations under strong positive selection, most of which are located in the capsid proteins and the 5’ noncoding region. By comparing viral diversity across household transmission pairs, we find that mOPV2 experiences a narrow transmission bottleneck which may limit the spread of mutations that are strongly selected within hosts. These results connect the within-host selection of mutations with the dynamics of viral transmission and enhance our understanding of cVDPV evolution.

## Results

We used high depth of coverage sequencing on the Illumina platform to define the within-host diversity of mOPV2 in samples from vaccinated individuals and their household contacts (29). These samples were collected as part of a cluster-randomized trial of OPV in the rural Matlab region, where the International Centre for Diarrheal Disease Research, Bangladesh (icddr,b) has conducted demographic and public health research since the 1960s (30). The trial assessed the impact of tOPV withdrawal on OPV community transmission by randomizing 67 villages to three different vaccination schedules: tOPV, bOPV followed by one dose of IPV, and bOPV followed by two doses of IPV. The trial then implemented a coordinated mOPV2 vaccination campaign over the course of one week, targeting 40% of children under 5 years of age. Shedding of OPV types 1-3 from 800 individuals across the three arms was monitored by quantitative RT-PCR of weekly stool samples. Transmission was measured by monitoring stool samples in household contacts of mOPV2 recipients. We selected 497 specimens from the vaccination campaign period for genome amplification and sequencing, prioritizing those with a stool viral load > 10^6^ copies/gram.

### Sample sequencing and assessment of genome coverage

We sequenced 416 samples from 219 mOPV2 recipients and 81 samples from 52 household contacts (Figure 1A). We amplified poliovirus genomes as four overlapping RT-PCR amplicons using degenerate primers that recognize all three OPV serotypes (Supplementary Table 1). We performed separate RT-PCR reactions for each segment and pooled them prior to library preparation (see Materials and Methods). Given that low viral titer influences the accuracy of within-host variant identification (31), we amplified and sequenced samples with an OPV2 viral load between 9 × 10^5^ copies/gram and 4.5 × 10^7^ copies/gram in duplicate. These cutoffs are based on the distribution of viral loads across the cohort and our empirically defined viral load cut-offs for influenza virus (32). We amplified and sequenced several samples below 9 × 10^5^ copies/gram that were collected from household contacts.

**Figure 1.**
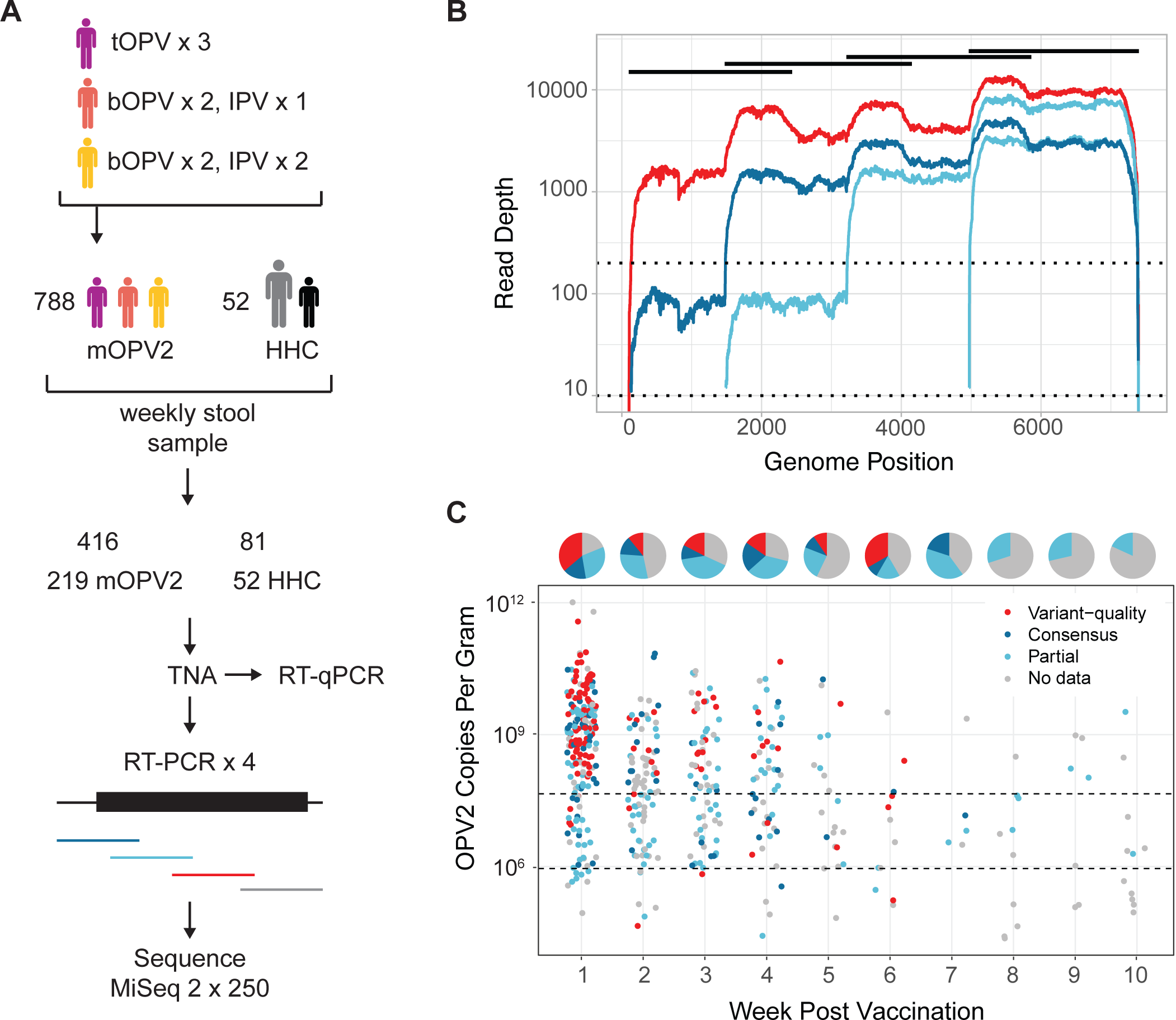
Overview of study and sequence data. (A) Schematic of study design and sample processing. The clinical trial had three arms with lead-up vaccination as indicated. tOPV = trivalent OPV, bOPV = bivalent OPV, IPV = inactivated polio vaccine. All individuals (n=788) then received mOPV2. Stool samples were collected weekly from mOPV2 recipients and household contacts (HHC). Only 52 household contacts had detectable shedding of OPV2. Total nucleic acid (TNA) was extracted from stools. Poliovirus genomes were amplified from each sample as four overlapping RT-PCR amplicons. For each sample, these amplicons were pooled and prepared for sequencing. (B) Line graph of sequencing coverage of four selected samples in three coverage groups. Log10 of coverage depth on the y-axis and genome position on the x-axis. One variant-quality sample shown in red, one consensus-quality sample shown in dark blue, and two partial-genome samples shown in light blue. Amplicons are shown as black bars (top). Dotted lines show cutoffs at 200x and 10x used for defining coverage groups. (C) Coverage groups of samples sequenced in this study. Each sample is shown as a point with OPV2 copies per gram of stool on the y-axis and week post-vaccination on the x-axis. Pie charts above each week indicate proportion of samples with variant-quality data (red), consensus quality data (dark blue), partial genome sequence data (light blue), no data (grey). Region in between the dotted lines shows the samples that were sequenced in duplicate.

Depth of coverage across the OPV2 genome for a given sample was uneven (Figure 1B). The 3’ end generally exhibited coverage at least an order of magnitude greater than the 5’ end, which contains the highly structured IRES (33). Three hundred twenty-seven samples had greater than 10x coverage of at least one of the four amplicons (Figure 1C, light blue points), and 179 samples had greater than 10x coverage across the whole genome (Figure 1C, dark blue points). We identified 111 samples with > 200x coverage (Figure 1C, red points), 81 samples with > 500x coverage, and 48 samples with > 1000x coverage across the genome, based on averages across a 50-bp sliding window. The majority of samples that yielded at least partial OPV2 genome coverage were collected in the first two months following mOPV2 vaccination (Figure 1C), which is consistent with the known shedding duration of Sabin type 2 poliovirus infections (34). Most individuals were represented by only one sample, although a subset of individuals had multiple longitudinal samples with at least partial genome data (Supplementary Figure 1).

### Empiric evaluation of variant calling criteria

We benchmarked the accuracy of our variant calling criteria for iSNV identification by sequencing defined mixtures of WPV 1 (Mahoney strain) and OPV1 in stool-derived total nucleic acid (TNA) with viral genome concentrations ranging from 9 × 10^4^ copies/gram to 4.5 × 10^7^ copies/gram of stool. These concentrations were tailored to match those of the sequenced clinical samples. We then calculated the sensitivity and specificity of iSNV identification at various thresholds of input concentration, sequencing coverage, and iSNV frequency (Table 1). At a coverage depth of 200x, we reliably identified the expected single nucleotide variants at 5% frequency with 95% sensitivity at all genome copy inputs. However, at this coverage level, sensitivity was weaker for low frequency variants. We found that the number of false positives was low when the viral load was greater than 4.5 × 10^7^ genome copies/gram. Specificity declined at viral loads below this cutoff, with a false positive rate of ∼1% at all coverage levels.

**Table 1:** Validation of within-host variant identification by sequencing mock populations.

(1) replicate, $4.5 \times 10^4$ copies/ $\mu\text{L}^a$ (2) replicates, $9 \times 10^3$ copies/ $\mu\text{L}^a$
| Coverage | Frequency | Sensitivity | Specificity | FP<sup>b</sup> | Sensitivity | Specificity | FP<sup>b</sup> |
| --- | --- | --- | --- | --- | --- | --- | --- |
| **200x** | 10% | 1 | 1 | 0 | 1 | 1 | 0 |
|  | 5% | 1 | 1 | 0 | 0.94 | 1 | 0 |
|  | 2% | 0.6 | 1 | 0 | 0.49 | 1 | 0 |
|  | 1% | 0.17 | 1 | 0 | 0.06 | 1 | 0 |
| **500x** | 10% | 1 | 1 | 0 | 1 | 1 | 0 |
|  | 5% | 1 | 1 | 0 | 1 | 1 | 0 |
|  | 2% | 0.91 | 1 | 0 | 0.74 | 1 | 0 |
|  | 1% | 0.54 | 1 | 0 | 0.4 | 1 | 0 |
| **1000x** | 10% | 1 | 0.9999 | 1 | 1 | 1 | 0 |
|  | 5% | 1 | 1 | 0 | 1 | 1 | 0 |
|  | 2% | 1 | 1 | 0 | 0.97 | 0.9999 | 1 |
|  | 1% | 0.91 | 1 | 0 | 0.69 | 1 | 0 |

(2) replicates, $9 \times 10^2$ copies/ $\mu\text{L}^a$ (1) replicate, $9 \times 10^2$ copies/ $\mu\text{L}^a$
| Coverage | Frequency | Sensitivity | Specificity | FP<sup>b</sup> | Sensitivity | Specificity | FP<sup>b</sup> |
| --- | --- | --- | --- | --- | --- | --- | --- |
| **200x** | 10% | 0.89 | 1 | 0 | 1 | 0.9995 | 7 |
|  | 5% | 0.83 | 1 | 0 | 0.80 | 0.9994 | 9 |
|  | 2% | 0.4 | 1 | 0 | 0.46 | 0.9990 | 14 |
|  | 1% | 0 | 1 | 0 | 0.31 | 0.9997 | 4 |
| **500x** | 10% | 0.97 | 1 | 0 | 1 | 0.9991 | 13 |
|  | 5% | 0.97 | 1 | 0 | 0.97 | 0.9985 | 21 |
|  | 2% | 0.49 | 1 | 0 | 0.51 | 0.9984 | 23 |
|  | 1% | 0.03 | 1 | 0 | 0.51 | 0.9992 | 12 |
| **1000x** | 10% | 1 | 1 | 0 | 1 | 0.9987 | 19 |
|  | 5% | 1 | 1 | 0 | 0.97 | 0.9974 | 37 |
|  | 2% | 0.63 | 1 | 0 | 0.91 | 0.9985 | 22 |
|  | 1% | 0.03 | 0.9999 | 2 | 0.03 | 0.9989 | 16 |

<sup>a</sup> Copies/ $\mu\text{L}$ is 1000-fold lower than copies/gram of stool.
<sup>b</sup> Number of identified false positives.

While some of these false positives can be filtered with various criteria, such as base and mapping quality, performing technical replicates proved to be the most effective approach for removing false positive variants. When considering only variants that were identified in both sequencing replicates, specificity dramatically improved even at low viral loads. Overall, these data validate our approach for poliovirus sequencing and demonstrate high sensitivity and specificity for variants above 5% frequency in the majority of sequenced samples. Therefore, we identified iSNV above a frequency threshold of > 5% in 111 samples that had > 200x coverage by sliding window across the genome, denoted here as “variant-quality.” For analyses of variants at particular genome positions, we included samples with > 200x mean coverage in a 50-bp window containing the site of interest.

### Signatures of selection within hosts

We characterized within-host diversity in 101 variant-quality samples from mOPV2 recipients. Minor iSNV (< 50% frequency) were dispersed across the genome in both the 5’ noncoding region and the polyprotein, with greater numbers of variants at lower frequencies (Figure 2A). Each sample contained a median of 9 minor iSNV with an interquartile range of 6-15. There were more minor iSNV per sample with greater time since vaccination (Figure 2B). This association remained significant even after we controlled for the time-varying factor of viral load, which can affect iSNV identification (p < 0.001, multiple linear model). Our estimates of minor iSNV frequency were consistent when compared between technical replicates of 11 variant-quality samples (adjusted r-squared = 0.763, Supplementary Figure 2A). While previous work has shown that the measured frequency of a variant can be affected by mutations in the primer binding sites (35), we were unable to distinguish this effect from the overall error in our frequency measurements. We identified a greater proportion of synonymous iSNV relative to nonsynonymous iSNV (ratio 0.43, Figure 2C). However, in the VP1 capsid subunit, there was an enrichment of nonsynonymous minor iSNV compared to other protein coding regions (Figure 2D). We calculated the dN/dS ratio with samples from mOPV2 recipients that had full consensus sequences across the polyprotein (n = 157). VP1 had the highest dN/dS ratio compared to the rest of the protein coding regions (Supplementary Table 2).

**Figure 2.**
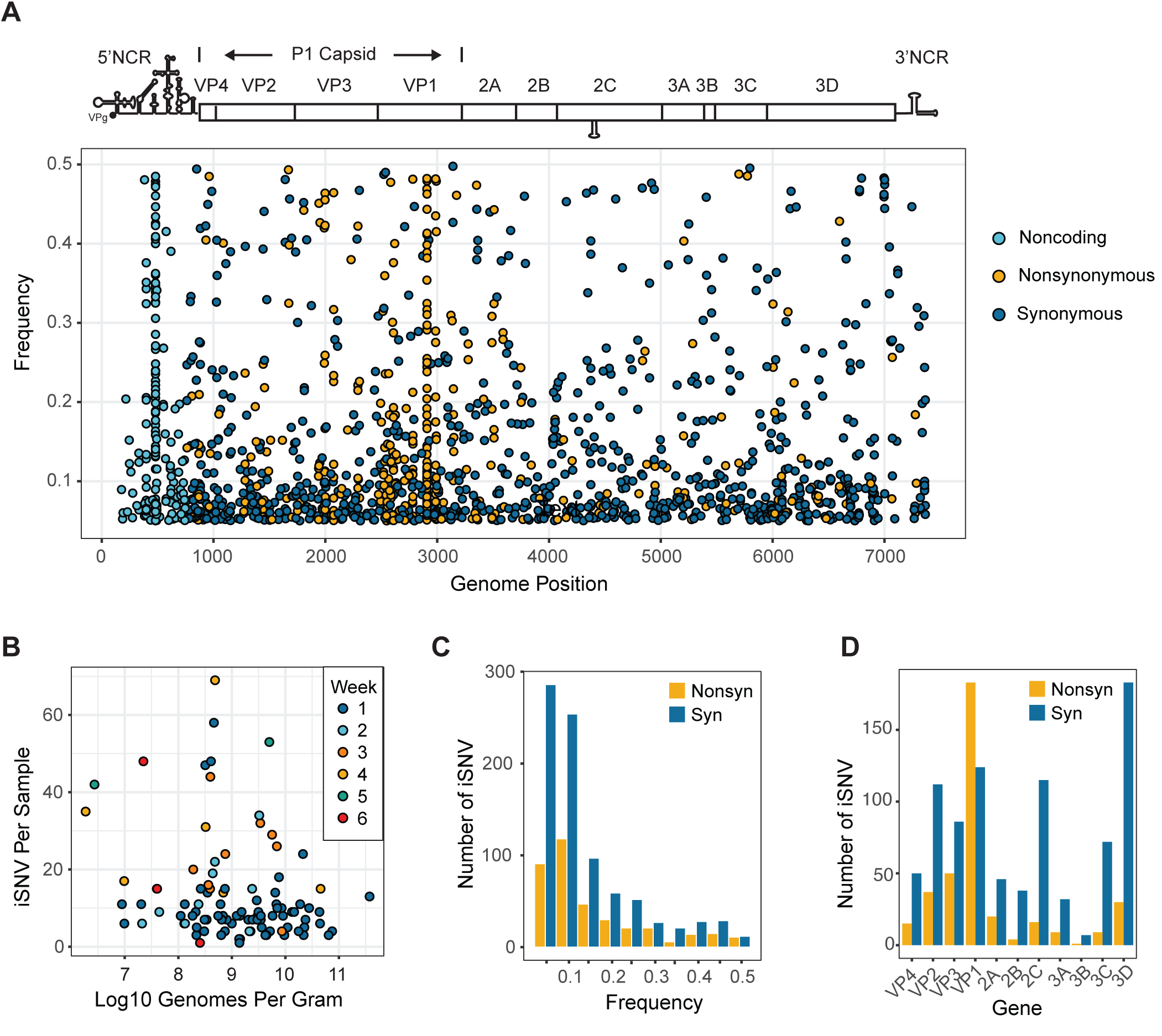
Within-host diversity in 101 variant-quality samples from mOPV2 vaccine recipients. (A) Minor iSNV shown as points, with frequency on the y-axis and genome position on the x-axis. Non-coding iSNV are shown in light blue, non-synonymous iSNV in yellow, and synonymous iSNV in dark blue. (B) Number of minor iSNV (y-axis) versus log_10_ of genome copies per gram of stool (x-axis). Color of each point is shown by the week post-vaccination of sample collection. (C) Histogram of minor iSNV in polyprotein by frequency with bin width of 0.05. Non-synonymous iSNV are shown in yellow, and synonymous iSNV in dark blue. (D) Histogram of minor iSNV by protein coding region in the polyprotein. Non-synonymous iSNV are shown in yellow, and synonymous iSNV in dark blue.

### Positive selection of gatekeeper mutations

The dN/dS ratio is an imperfect metric for detecting selection, particularly within hosts, and it is unable to identify positive selection of mutations in noncoding regions (36). While changes in frequency of viral variants can be caused by multiple evolutionary forces, observing the same mutation arise in independent viral populations is suggestive of positive selection (37, 38). We therefore analyzed the mutational dynamics at three positions that are major attenuating sites – positions 481 and 398 in the 5’ noncoding region and codon 143 of VP1 (nucleotide positions 2908-2910). We used our time-series data to directly measure the frequency changes of the gatekeeper mutations in vaccine recipients. All three mutations were present in several individuals within the first week of vaccination. A cross-sectional analysis of mutation frequency as a function of time demonstrated fixation of A481G within 2-3 weeks and U2909C in about 5 weeks, although there was substantial interindividual variability (Figure 3A and 3B). While A481G reached consensus in 11 of 14 samples by week 2, U2909C reached consensus in only 5 of 22 samples by week 3. VP1-143 most frequently reverted from isoleucine to threonine, but several other alternative residues were present (Figure 3C). Data from individuals with more than one sequenced sample demonstrated a rapid increase in frequency. Although mutation frequency occasionally decreased, presumably due to stochastic effects, these mutations increased in nearly all individuals (Figure 3B and 3D). We applied a beta regression model to estimate the time to fixation in the population. For mutations A481G, U2909C, and U398C, the model predicts a frequency of > 0.5 at weeks 2, 5, and 12, respectively, and > 0.95 by weeks 6, 13, and 46, respectively (Supplementary Figure 3). While we had more data from individuals in the bOPV/IPV study arms, each mutation rose in frequency over the same time interval regardless of vaccination history. This suggests that the selection for these mutations is not substantially driven by the presence of mucosal immunity generated by tOPV (Supplementary Figure 3). Overall, our time-series data on mOPV2 recipients shows rapid fixation of mutations at key attenuating sites in the first several weeks post-vaccination.

**Figure 3.**
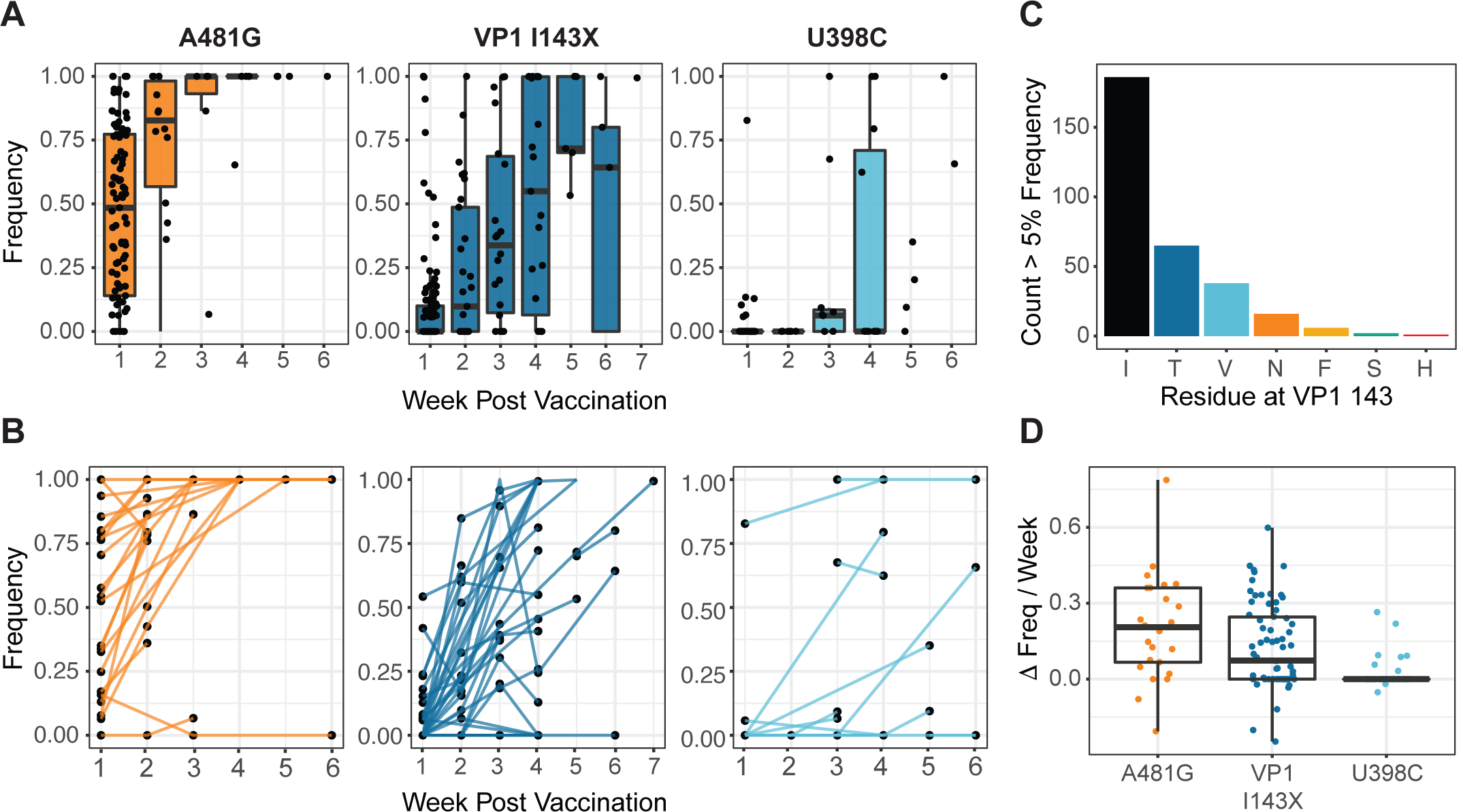
Selection of gatekeeper mutations in vaccine recipients. (A) Frequency of A481G, VP1-143X, and U398C by time from vaccination. Each point represents one sample, and boxplots are shown for weeks with five or more data points. Boxplots represent the median and 25^th^ and 75^th^ percentiles, with whiskers extending to the most extreme point within the range of the median ± 1.5 times the interquartile range. (B) Frequency of A481G, VP1-143X, and U398C by time from vaccination. Each point represents one sample, with lines connecting samples from the same individual. (C) Barplot showing number of samples with the indicated residues present at a frequency of 5% or above at VP1-143. (D) Change in frequency per week of three gatekeeper mutations prior to reaching fixation. Boxplots represent the median and 25^th^ and 75^th^ percentiles, with whiskers extending to the most extreme point within the range of the median ± 1.5 times the interquartile range.

### Additional sites with positive selection

We next identified non-gatekeeper mutations that occurred independently across mOPV2 recipients. We restricted our analysis to 83 individuals who received mOPV2 and for whom we had at least one variant-quality sample. While most mutations were unique to a given viral population, a large number of mutations were present at ≥ 5% frequency in two or more individuals (Figure 4A). We performed a permutation test to quantitatively assess whether this distribution could occur due to chance alone. We drew random sites across the genome and tallied the number of sites shared across multiple individuals (see Materials and Methods). We observed more shared mutations than would be expected by chance for mutations in three or more individuals (Figure 4A). This result is robust to the assumption that all genome sites can be mutated; reducing the fraction of sites available for mutation did not affect statistical significance until the fraction dropped below 50% (Supplementary Figure 4).

**Figure 4.**
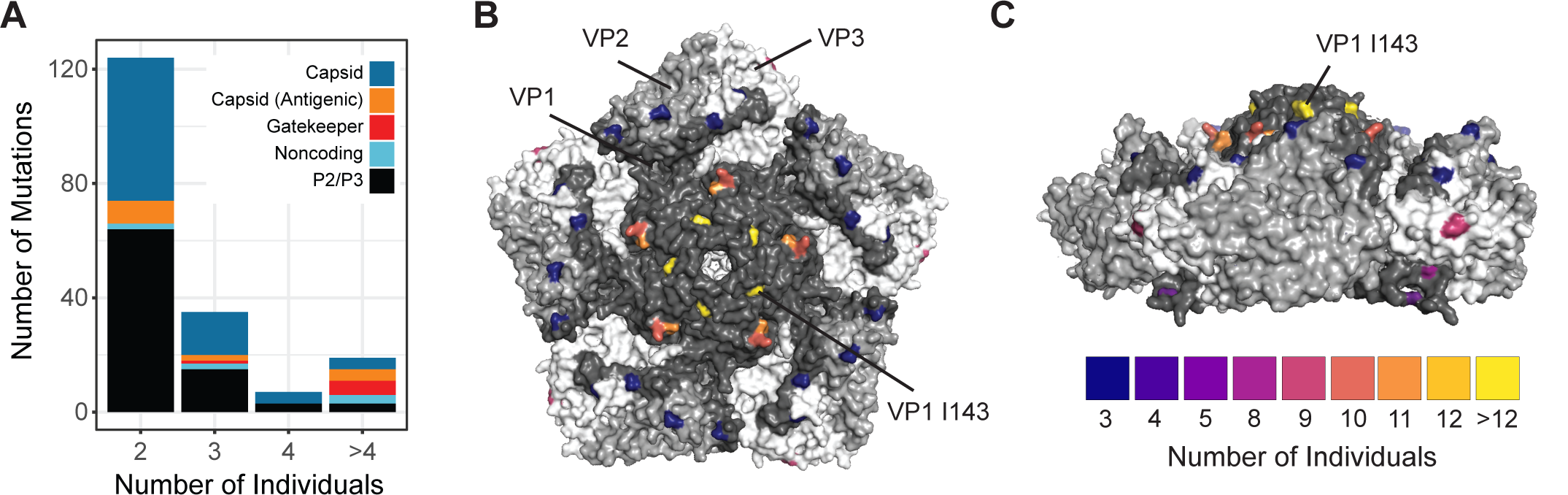
Mutations arising in multiple mOPV2 vaccine recipients. (A) Stacked barplot of the number of mutations identified (y-axis) by the number of individuals with each mutation (x-axis). Mutations in ≥ 3 individuals were statistically significant by permutation test, see text. Colors show the category of each mutation. (B) Structure of type 2 poliovirus capsid pentamer (PDB: 1EAH) and side view (C). Highlighted residues are color-coded by number of mOPV2 vaccine recipients with non-synonymous substitutions at that amino acid site.

Excluding the gatekeeper mutations, we found 19 mutations that were present in four or more individuals (Table 2). Two mutations, G491A and G619U, were located in the IRES. Five mutations were located outside the capsid (P1) in polyprotein regions P2 and P3. The capsid proteins (VP1-4) were highly represented (12 of 19 mutations), with four mutations encoding nonsynonymous mutations in known antigenic sites. Mutation A2986G encodes a nonsynonymous substitution, VP1-K169E, in antigenic region NAg1. Two more mutations in NAg1, G2782A and C2783A, encode nonsynonymous changes at VP1-101 (A101T and A101D, respectively). Mutation A1997G, found in 8 of 83 individuals, encodes VP3-H77R in antigenic region NAg3b. This mutation was identified in a previous phylogenetic study as having intermediate evidence for positive selection across cVDPV lineages (9). Our host-level data show that this mutation is indeed under strong positive selection. We did not detect U2523C, C2006A, U1376A, and U3320A, which are predicted to occur > 2 months after vaccination (9).

**Table 2:** Mutations identified in multiple individuals.

| Mutation<sup>a</sup> | Individuals<sup>b</sup> | Group | Type<sup>c</sup> | Region | Fraction of cVDPV<sup>d</sup> |
| --- | --- | --- | --- | --- | --- |
| A481G | 72 | Gatekeeper | Noncoding | 5' UTR | 1 |
| U2909C | 25 | Gatekeeper | NS | VP1 | 0.75 |
| A2908G | 19 | Gatekeeper | NS | VP1 | 0.03 |
| U398C | 16 | Gatekeeper | Noncoding | 5' UTR | 0.94 |
| A2074G | 12 | Capsid | NS | VP3 | 0.01 |
| A2992G | 11 | Capsid | NS | VP1 | 0.03 |
| A2986G | 10 | Antigenic | NS | VP1 | 0.01 |
| G6084U | 10 | 3D | S | 3D | 0.03 |
| A1997G | 8 | Antigenic | NS | VP3 | 0.05 |
| U2909A | 8 | Gatekeeper | NS | VP1 | 0.03 |
| U882C | 8 | Capsid | S | VP4 | 0.02 |
| G2782A | 7 | Antigenic | NS | VP1 | 0.01 |
| G491A | 7 | Noncoding | Noncoding | 5' UTR | 0.14 |
| G619U | 7 | Noncoding | Noncoding | 5' UTR | 0 |
| C2609U | 6 | Capsid | NS | VP1 | 0.01 |
| C2783A | 5 | Antigenic | NS | VP1 | 0 |
| U4374C | 5 | 2C | S | 2C | 0.55 |
| A3490G | 4 | 2A | NS | 2A | 0.02 |
| C2291U | 4 | Capsid | NS | VP3 | 0.01 |
| C2580U | 4 | Capsid | S | VP1 | 0.1 |
| G1282A | 4 | Capsid | NS | VP2 | 0 |
| U1641C | 4 | Capsid | S | VP2 | 0.68 |
| U5811A | 4 | 3C | S | 3C | 0.14 |
| U6693A | 4 | 3D | S | 3D | 0.31 |

<sup>a</sup> Mutation presented as base in OPV2, position in OPV2 reference genome, and base in samples.
<sup>b</sup> Number of individuals with each mutation present at a frequency of 5% or greater. The total number of individuals analyzed is 83.
<sup>c</sup> Nonsynonymous (NS) or synonymous (S) mutations relative to the OPV2 reference genome.
<sup>d</sup> Fraction of cVDPV genomes with each mutation.

We also examined independent capsid mutations at the codon level, such that distinct mutations at the same amino acid site were included. We identified 19 amino acid sites at which three or more individuals exhibited nonsynonymous substitutions at a frequency of > 5%. Many of these sites mapped to the surface of the type 2 poliovirus capsid (Figure 4B). In addition to the amino acid sites specified above, this analysis revealed four more antigenic sites with parallel non-synonymous substitutions: VP3-58 (T58I and T58A) and VP1-T291A in NAg3a, VP2-N72D in NAg3b, and VP1-S222P in NAg2. Although dN/dS analysis is often not sensitive enough to identify positive selection at specific sites over short time scales, amino acid sites VP1-143 and VP1-101 had a dN/dS ratio greater than 1 based on analysis of consensus genomes (Pr(*ω* > 1) > 0.95, Bayes empirical Bayes method, Supplementary Table 2). Together, our results demonstrate rapid positive selection of mutations in the IRES and exposed sites in the OPV2 capsid.

We sought to determine whether mutations selected early in OPV2 evolution persist in genomes of neurovirulent cVDPV. We queried alignments of cVDPV from cases of acute flaccid paralysis for the presence of the mutations identified here (Table 2). As expected, the gatekeeper mutations were reliably detected in nearly all cVDPV genomes. Of the other 19 mutations identified in our analysis, we found 16 in at least 1% of cVDPV genomes queried. We detected some mutations, including G491A, C2580U, and U1641C, in at least 10% of cVDPV genomes. While it is unknown how these mutations may contribute to the development of cVDPV outbreaks, our data show that mutations that recur in divergent cVDPV lineages can be identified very early in OPV2 evolution.

### Estimation of the transmission bottleneck

Transmission bottlenecks influence the rate of viral adaptation and are important for understanding viral evolution in host populations (39). The OPV vaccine trial included stool samples collected from the household contacts of mOPV2 recipients. Shedding in these household contacts allowed us to analyze the extent of viral diversity that is transmitted to new hosts. We identified four transmission pairs for which we had sequence data from both the donor and recipient collected within one week of each other (Supplementary Table 3); sequencing of more putative household pairs was thwarted by low viral loads, especially in household contacts. In each case, household transmission occurred within the first three weeks after vaccination, consistent with the known magnitude and duration of OPV shedding. We obtained variant-quality samples from donor and recipient of one pair; the rest had either consensus or partial genomes with regions of variant-quality coverage.

We compared within-host diversity across donors and recipients using only the genomic regions that had a depth of coverage sufficient for identification of within-host variants in both samples. There were few polymorphic sites shared across hosts in these household pairs (Figure 5A). While major variants (> 50% frequency) in the donor were usually found in the recipient, most minor variants were not found in the recipient, suggestive of a narrow transmission bottleneck.

**Figure 5.**
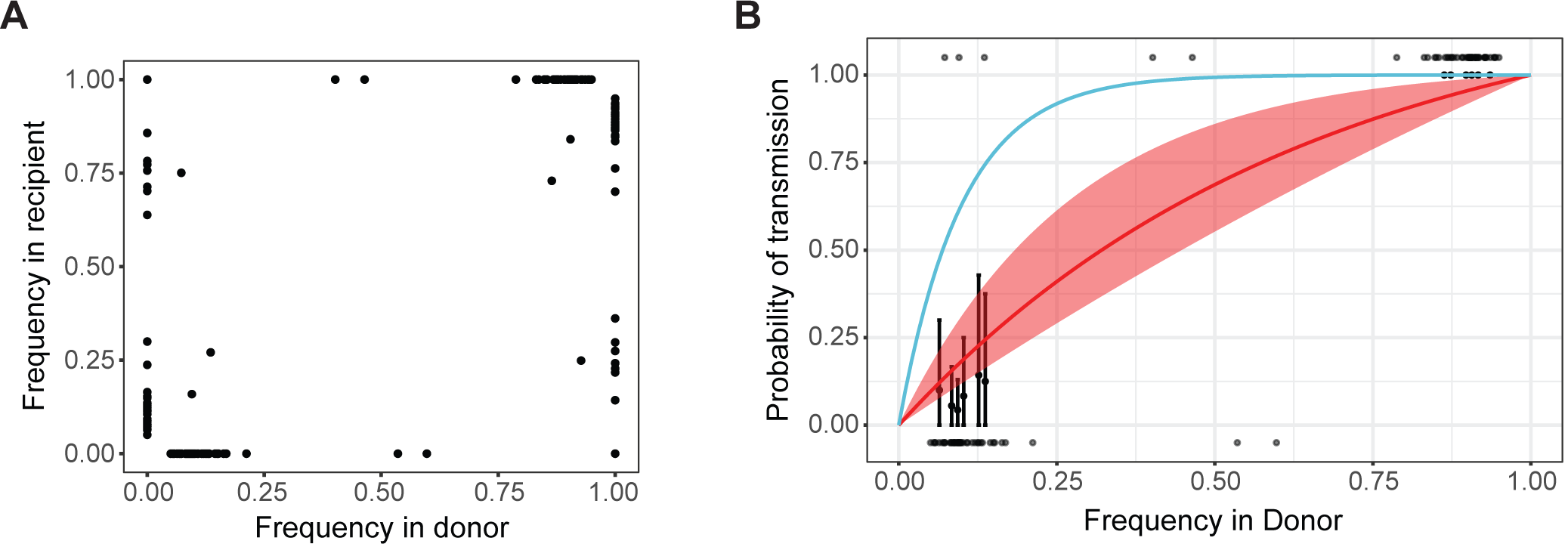
Shared viral diversity across transmission pairs and transmission bottleneck. (A) iSNV for four pairs of mOPV2 recipients and their household contacts. Each iSNV is plotted as a pointwith its frequency in the recipient (y-axis) versus its frequency in the donor (x-axis). (B) Presence-absence bottleneck model fit compared with data. Frequency of donor iSNV on the x-axis and probability of transmission on the y-axis. Transmitted iSNV are shown along the top of the plot and non-transmitted iSNV are shown along the bottom. The red line shows the probability of transmission as a function of donor frequency given the mean bottleneck estimate, with a 95% confidence interval shown by the shaded area. The blue line shows the probability of transmission given a bottleneck size of 10. The black points on the graph represent the probability of transmission from the measured iSNV using a sliding window of 3% width and a step size of 1.5%.

We applied two models to quantify the effective genetic bottleneck at transmission (32, 40). The presence-absence model asks whether a polymorphism in the donor is present or absent in the recipient, assuming perfect detection. Here, transmission is modeled as a random sampling process in which the probability of transmission is a function of the mutation frequency in the donor and the size of the transmission bottleneck. We used maximum likelihood optimization to find the bottleneck size distribution that best fit the data, assuming that the bottlenecks follow a zero-truncated Poisson distribution. Under the presence-absence model, the mean bottleneck size was 1.98 (lambda = 1.57, 95% confidence interval 0.43 – 3.63), indicating that most bottlenecks are 2 and 95% of bottlenecks are less than 4 (Figure 5B). We also applied a beta-binomial model, which incorporates the sensitivity of detecting variants in the recipient and allows for time-dependent stochastic loss of variants. The beta-binomial model yielded a mean bottleneck size of 2.11 (lambda = 1.74). The model fit was not significantly better than the presence-absence model (AIC 42.1 for presence-absence vs. 39.7 for beta-binomial), indicating that the loss of sensitivity in the recipient might not be an important factor in these models. We also estimated bottlenecks for each pair individually with both models (Supplementary Table 4). The two models produced the same estimates for each pair, although the beta-binomial model resulted in slightly larger confidence intervals. These results suggest that few genetically distinct OPV2 genomes transmit and persist in new hosts.

These models assume that minor variants are transmitted independently. If this assumption is violated by linkage between minor variants, this could result in an inflated bottleneck estimate. Although our estimate is already low, we sought to assess the extent of linkage between minor variants in these data. We focused on the minor variants in the donor of pair 702, for which we had variant-quality data on both donor and recipient populations. From the 20 minor variants present in this donor, we found 18 pairs of minor variants for which we had reads that spanned the variant sites. For each minor variant, we found the fraction of reads that also contained the other minor variant in the pair. Out of the 18 variant pairs, only two had both minor iSNV present in greater than 50% of reads, indicating physical linkage (Supplementary Figure 5). None of the presumably linked variants were transmitted to the recipient host. These data indicate that while linkage between minor variants can and does occur, most of the variants modeled in these hosts are unlinked.

### A tight transmission bottleneck limits the spread of gatekeeper mutations

We sought to investigate whether a narrow transmission bottleneck would impact the transmission of mutations that are positively selected within vaccine recipients. Based on the estimated bottleneck size, we calculated the probability of transmission of each of the three gatekeeper mutations as a function of time, using the median frequency from mOPV2 recipients (Figure 6A). We then calculated the fraction of samples from transmission recipients that had each mutation present, regardless of whether we had sequence data from a donor population (Figure 6B). The fraction of transmission samples with each mutation is consistent with the calculated probability of transmission over time given the size of the bottleneck. We identified A481G in most transmission samples, consistent with its rapid fixation. However, few samples contained U2909C and no samples contained U398C. The majority of transmission events occurred within the first two weeks after the vaccination campaign, prior to when most vaccine recipients acquired U2909C and U398C (29). These data suggest that the three gatekeeper mutations were not preferentially transmitted; instead, they suggest that mutations must rise to an appreciable frequency early enough within a donor population to be frequently transmitted through a narrow bottleneck.

**Figure 6.**
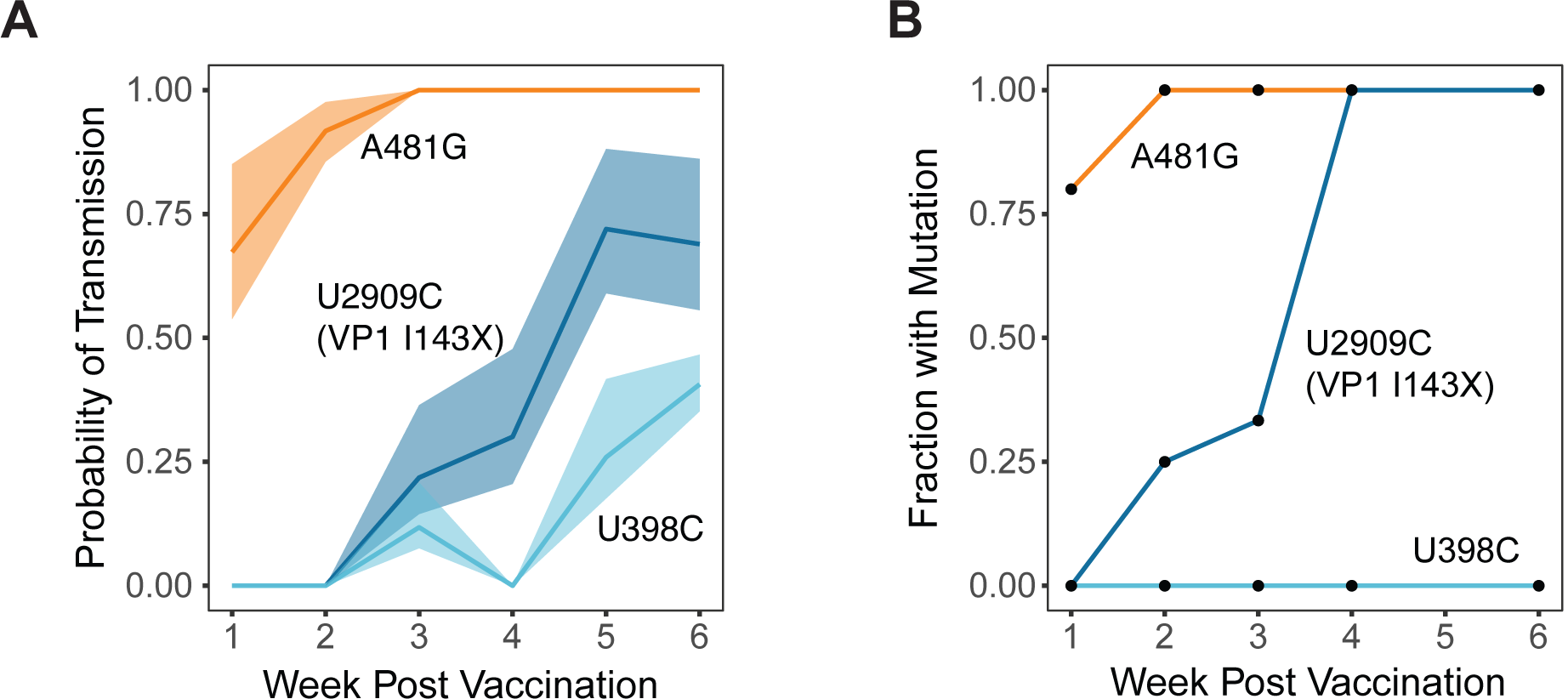
Impact of a tight bottleneck on transmission of gatekeeper mutations. (A) The probability of transmission of each gatekeeper mutation calculated from the median frequency over time in the mOPV2 recipients given the estimated bottleneck. The shaded areas represent 95% confidence intervals based on the model fit. (B) The fraction of each gatekeeper mutation present above a frequency of 5% in samples from household contacts as a function of time since the vaccination campaign.

## Discussion

We used whole genome deep sequencing to define the within-host evolutionary dynamics of OPV2 in a clinical trial in Matlab, Bangladesh (29). The vaccine trial enabled analyses of longitudinal samples from a defined and synchronized point of mOPV2 vaccination and household transmission in a community with high enteric pathogen burden and vaccine coverage. These results provide a rare window into the evolutionary dynamics that occur in the first weeks following vaccination. Similar to other RNA viruses, we identified strong purifying selection across the poliovirus genome within hosts (32). However, in stark contrast to other viruses, we found evidence for strong within-host positive selection across multiple sites. Although high population immunity in Bangladesh limited the number of transmission samples available from the trial, we were able to quantify the transmission of key reversion mutations and estimate a tight bottleneck in this setting. Our findings enhance our knowledge on the within-host and transmission dynamics of polioviruses in relation to the development of cVDPV.

We find that positive selection is remarkably strong within vaccine recipients, with a magnitude that is seldom found in similar studies of within-host evolution of acute RNA viruses. We and others have rarely found strong selection for mutations at the within-host level, even for mutations that should have beneficial effects (41, 42). The within-host evolution of several arboviruses is characterized by purifying selection and a large effect of stochastic genetic drift (43, 44). In household cohort studies of influenza virus infection, we have found little evidence for positive selection within the span of a single infection (32, 45), and iSNV are rarely observed in more than one individual. In contrast, in this cohort we identified 24 mutations that were identified in four or more individuals. Although comparisons to other viruses are complicated by differences in duration of infection, genome structure, and other factors, the extent of parallel evolution in OPV at this scale is remarkable. Whereas wild polioviruses and other endemic RNA viruses may already exist near local fitness peaks, OPV is significantly attenuated and is under intense pressure to climb the fitness landscape by accessing available high-impact mutations (9). OPV is also unique in that each population starts from the same founder genetic sequence, making parallel trajectories more likely to occur (38).

Outside of the three gatekeeper mutations, we find that there are multiple additional mutations under selection early in OPV2 evolution that reflect re-adaptation to the human host. There are several potential reasons why our study revealed mutations that have not been previously identified in cell culture or phylogenetic studies. While cell culture and animal models can be a helpful proxy for inferring selective pressures on viruses (17), they do not always capture the direction and magnitude of evolutionary forces in natural hosts. Similarly, phylogenetic studies have yielded important insights into the evolution and epidemiology of cVDPV, but may not be able to infer selective advantage of mutations with weaker effects due to limitations in sampling or statistical power. Whereas, previous phylogenetic work on cVDPV2 found only limited evidence for positive selection at VP3-77 and VP1-222, they appear to be strongly selected within hosts (9, 11). Finally, it is important to recognize that phylogenetic studies may differ in the fitness effects they reveal due to differences in time and scale. Our results primarily reflect within-host selection observed through parallel evolution among vaccine recipients sampled longitudinally whereas previous phylogenetic analyses are based on shared variation among surveillance samples collected from many people over time and linked by sustained transmission. While we found that 16 of the 19 positively-selected, non-gatekeeper variants recur in cVDPV lineages, they do not routinely fix. Some mutations might be repeatedly selected within hosts and prove to be detrimental between them.

The underlying selective pressures and consequences of these mutations are unclear, but their locations suggest functional significance. Mutations in the IRES have been shown to affect viral protein translation and replicative capacity (46). Mutations in the capsid enable adaptation to replication at physiologic temperatures, and could provide increased structural stability or modulate receptor binding and viral entry (15, 47). Although some of the capsid sites identified here are recognized by neutralizing antibodies (11, 48), there is little evidence that these mutations lead to antigenic escape, as they do in influenza virus or HIV (49). Even highly diverged cVDPV strains with significant antigenic site evolution are still neutralized by serum from vaccinated individuals (11), and vaccination with OPV is used to control outbreaks of cVDPV (6). Rather than antigenic escape, we suggest that these mutations lead to improved within-host replication, and therefore greater shedding and transmission. Epidemiologic data suggest that at some unknown point in cVDPV evolution, OPV achieves a level of transmissibility that is similar to that of wild polioviruses (2, 34, 50). Selection for phenotypes related to this increase in transmissibility, like enteric replication and shedding, are likely the earliest pressures the virus faces (51). Not all capsid antigenic sites may be involved in this process. There can be frequent amino acid substitution at many antigenic sites in the OPV2 capsid, often reverting back to previous replacements, suggesting that some sites are more tolerant to mutation and evolve more by genetic drift than selection (11). However, our results indicate that a subset of these capsid sites experience positive selection and likely have functional effects related to improved replication and transmission within the human host.

Our identification of specific sites under positive selection has implications for genetic surveillance of VDPV. In VDPV isolates, the time since vaccine administration is estimated by molecular clock methods on VP1 sequence data (52). For OPV2, the threshold for calling a strain a VDPV – as opposed to OPV-like – is 0.6% divergence, or *≥* 6 nucleotide substitutions (53). Prior work has integrated the fixation rates of gatekeeper mutations into molecular clock models to refine estimates of the time between VDPV detection and initial vaccination (10). By accounting for these rapidly selected mutations, the authors inferred that type 2 VDPVs are younger than estimated based on neutral evolution alone. Here, we used our longitudinal data to determine the fine-scale dynamics of these three gatekeeper mutations and to characterize the variability that can manifest at the individual scale. Fixation rate estimates that are grounded in direct measurements are relevant for modeling efforts that rely on these parameters. In addition, we suggest that molecular clock models of VDPV might benefit from incorporation of the novel sites identified in this work in two ways. First, the rates of selection at these sites could be integrated into existing models of time since vaccine dosing for VDPVs. Second, these sites under putative selection could be excluded from neutral molecular clocks for VDPV divergence time estimation.

A surprising and important finding is that a tight bottleneck (1-4 distinct genomes) limited the transmission of within-host variants to new hosts. It is certainly possible that loss of variants in the recipient population could result in an under-estimation of the bottleneck. In this study, we were limited by the week-long interval between sample collection, which means that transmission could have occurred several days prior to sampling from the household contact. We also used a conservative frequency threshold of 5%, which may miss transmission of variants that remain at low frequencies across both hosts. However, it is unlikely that variants below 5% would be consistently transmitted while variants from 5-50% are not. Furthermore, the results of the beta-binomial model suggest that imperfect detection and stochastic loss did not have a large influence on the bottleneck estimate.

In support of the finding of a small bottleneck in this transmission setting, we have no evidence for a between-host advantage of the gatekeeper mutations despite strong evidence of within-host selection. If genomes with gatekeeper mutations are transmitted preferentially, we would have expected to see them present in more household contacts and at higher frequencies than observed here. The sample size in this study limits our ability to detect small effects, but the data generally do not support substantial preferential transmission in this study population. A narrow, non-selective bottleneck can explain the pattern of transmission to household contacts. In this scenario, mutations selected within vaccine recipients must rise above a threshold frequency prior to transmission in order to transit a narrow bottleneck. This would limit spread of minor iSNV that have not been selected quickly enough before finding a new host.

Of course, the gatekeeper mutations eventually fix in nearly all vaccine-derived lineages and can arise *de novo* in each subsequent host (9, 10). Population immunity and differences in fecal-oral exposure between Matlab, which has never experienced a cVDPV outbreak, and other settings where cVDPV outbreaks are more common may lead to important selective differences not observed here. In contrast to the lack of between-host selective effects in this study, late transmission events at the end of the duration of shedding when novel variant fractions are high, may have a larger impact on the spread of positively selected mutations. Transmission later in infections when viral load is low and to more distant community contacts was uncommon in this highly immune population (29). Furthermore, bottlenecks may be larger in populations with lower background immunity and higher fecal-oral exposure where naturally acquired doses are likely higher (34), which would allow positive selection among minor variants to act.

It is likely that newly developed live-attenuated polio vaccines will face the same underlying selection pressures as mOPV2. Our results suggest that once a beneficial mutation occurs on a highly attenuated OPV background, there is strong selection to drive it to fixation. Strategies that decrease the fitness benefit of any single mutation, like modifications to IRES domain V and codon deoptimization, may be effective at sufficiently prolonging the time to reversion (54, 55). However, modest decreases in mutation rate by introduction of high-fidelity RNA-dependent RNA polymerase modifications might have lesser impact, as the mutation rate is still orders of magnitude higher than in other organisms (56). In the setting of a virus starting from low fitness with a high mutation rate, whether a mutation achieves fixation or not may be more dependent on size of the fitness benefit rather than the waiting time for *de novo* generation of the mutation. This effect is illustrated by one novel OPV2 (nOPV2) design, which prevents A481G by modification of IRES domain V. In individuals receiving this nOPV2, VP1-143 and U398C still readily revert despite a high-fidelity 3D^pol^ (54). High-fidelity polymerase modifications themselves may not be stable, as seen by the reversion and compensation of a type 1 poliovirus fidelity mutant due to a fitness defect in cell culture (57). Our results also suggest that there are mutations other than the three gatekeepers that increase fitness and contribute to reversion. Monitoring the genetic changes of new vaccine designs in sufficiently large cohorts will be important for evaluation of the genetic stability at these additional sites.

## Materials and Methods

### Clinical trial information and ethics

The clinical trial, including all aspects of sample collection and viral load measurements, is described in full in Taniuchi et al. (29). The study was done according to the guidelines of the Declaration of Helsinki. The protocol for the clinical trial was approved by the Research Review Committee (RRC) and Ethical Review Committee (ERC) of the icddr,b and the Institutional Review Board of the University of Virginia. It is registered at ClinicalTrials.gov, number NCT02477046.

### Sample collection and viral load quantification

Briefly, stool samples were collected, placed at 4°C, and delivered to the icddr,b laboratory in Matlab within 6 hours. Samples were then aliquoted and stored at −80°C until shipment on dry ice to the icddr,b laboratories in Dhaka. Total nucleic acid (TNA) from approximately 200 grams of stool was extracted with the QIAamp Fast DNA Stool mini kit and OPV was detected and quantified by RT-qPCR with serotype specific primers. TNA samples were shipped on dry ice to the University of Virginia and stored at −80 °C until processing for sequencing.

### Primer design

We designed primers to amplify all three serotypes of OPV in four overlapping amplicons covering the poliovirus genome. We used PrimerDesign-M (58) to determine sites of conservation across the three poliovirus types and identify potential primer sequences, allowing for ambiguous bases. We selected primers such that the four segments overlapped by at least 500 bp and manually curated each primer to have similar melting temperatures. We empirically tested various primer candidates for amplification on type 1 and type 2 poliovirus RNA templates. The primers used for genome amplification are listed in Supplementary Table 1.

### Amplification and sequencing

We amplified viral cDNA in four amplicons using a two-step RT-PCR protocol. We performed reverse transcription using the SuperScript III First-Strand Synthesis System (ThermoFisher). Each reaction contained 1.13 μL of 50 ng/μL random hexamers, 0.37 μL oligo-dT, 1.5 μL 10 mM dNTP mix, 12 μL of template total nucleic acid from stool, 3 μL of 10x RT Buffer, 6 μL 25 mM MgCl2, 3 μL 0.1M DTT, 1.5 μL RNase Out, and 1.5 μL of SuperScript III RT enzyme. The mixture of template, primer, and dNTPs was heated at 75°C for 15 minutes to denature RNA secondary structure and placed directly on ice for > 1 minute. The enzyme and buffer mixture were then added on ice. The thermocycler protocol was: 25°C for 10 min, 50°C for 50 min, 85°C for 5 min, and hold at 4°C. The four overlapping segments were amplified by PCR with the primers in Supplementary Table 1. The PCR reactions were as follows: 10 μL 5x HF Buffer, 1 μL 10 mM dNTP mix, 0.25 μL 100 μM forward primer, 0.25 μL 100 μM reverse primer, 33 μL nuclease free water, 0.5 μL of Phusion DNA Polymerase (NEB), and 5 μL of template cDNA. The thermocycler protocol was: 98°C for 30 sec, 40 cycles of 98°C for 10 sec, 59.5°C for 30 sec, 72°C for 2 min, then 72°C for 2 min for final extension, and hold at 4°C. The four segments for each sample were pooled in equal volumes (18 μL of each segment for a total pooled volume of 72 μL). Pooled amplicons were purified with Agencourt AMPure XP magnetic beads, using 1.8X volume of beads (129.6 μL of beads for 72 μL pooled PCR product). The sample was eluted into 40 μL of nuclease-free water. The purified PCR products were quantitated by Quant-iT PicoGreen dsDNA High Sensitivity Assay. A limited number of PCR products were spot-checked by gel electrophoresis. A plasmid control was prepared by applying the PCR protocol to a template of OPV2 in a plasmid. The sequence of the plasmid was determined by Sanger sequencing and was identical to the OPV2 GenBank reference (AY184220.1). One plasmid control was included in each pooled library, beginning at the library preparation stage, to account for sequencing errors and batch effects. Samples between 9 × 10^5^ copies/gram and 4.5 × 10^7^ copies/gram of OPV2 were amplified and sequenced in duplicate to improve the specificity of within-host variant identification. Libraries were prepared for Illumina sequencing with the Nextera DNA Flex Library Preparation kit according to the manufacturer’s instructions, using Nextera DNA CD Indexes (96 samples). Eight pooled libraries were prepared in total and sequenced on an Illumina MiSeq (2×250 reads, v2 chemistry).

### Benchmarking of variant identification with a mock population

To determine the sensitivity and specificity of variant identification, we sequenced mock populations of a mixture of two viruses using the protocol described above. The viruses used were wild-type Mahoney type 1 poliovirus and the type 1 OPV strain, which differ by 66 mutations within the amplified regions. The plasmids containing each virus were confirmed by Sanger sequencing. We generated passage 1 viral stocks for each virus in HeLa cells and mixed them at equal concentrations at 0%, 1%, 2%, 5%, 10%, and 100% WT in OPV1. Concentrations of poliovirus genome copies of each virus stock were determined by RT-qPCR and diluted to concentrations of 4.5 × 10^4^ copies/μL, 9 × 10^3^ copies/μL, 9 × 10^2^ copies/μL, and 9 × 10^1^ copies/μL (copies/gram is related to copies/μL by a factor of 10^3^). To simulate the complex mixture of nucleic acids present in our samples, we performed the dilutions of viral populations in total nucleic acid extracted from stool from deidentified human donors (a gift of Pat Schloss, University of Michigan). Total RNA was extracted from each unique mock population and amplified by the protocol described in the section above (*Amplification and sequencing*). A plasmid control was generated from the OPV1 plasmid in the same way as described above. The pooled library was generated using the Nextera DNA Flex Library Preparation Kit and sequenced on an Illumina MiSeq (2×250 reads, v2 chemistry), including the OPV1 plasmid control to account for batch effects and errors. To more carefully estimate the sensitivity of variant identification at various coverage levels, we performed random down-sampling of the mapped reads to approximately 1000x, 500x, and 200x coverage evenly across the genome.

Then we identified within-host variants using an analytic pipeline previously used for influenza viruses (31, 32), which depends on a clonal plasmid control to account for batch effects and local errors in Illumina sequencing.

### Identification of consensus sequences and phylogenetic analysis

Sequencing adapters were removed with cutadapt (59) and reads were aligned to all three OPV reference genomes (AY184220.1, AY184221.1, V01150.1) using bowtie2 (60) with the –very-sensitive option. Duplicate reads were removed with Picard and samtools (61). Consensus bases were identified at sites with 10x coverage or greater. For samples sequenced in duplicate, the replicate with the higher coverage at a given site was used to assign the consensus base. Each biological sample was assigned to a group based on depth and evenness of coverage across the OPV2 reference. Mean coverage was calculated across non-overlapping 50 bp bins across the genome region amplified by the four amplicon segments. Samples were considered variant-quality if they had an average coverage greater than 200x in every bin. Samples that had coverage of 10x or greater at every site were considered consensus-quality. We aligned the consensus sequences with the OPV2 reference with the MUSCLE algorithm (62). For the dN/dS analysis, we used the PAML software version 4.8 (63). For gene-wise dN/dS analysis, we calculated a single value for omega across each gene with codeml using model M0. To identify sites with evidence of positive selection, we compared the likelihood of models M2 vs M1 with a chi-squared test and identified sites with omega greater than 1 with the Bayes empirical Bayes method.

### Identification of within-host variants

We identified within-host variants in any 50 bp window with greater than 200x mean coverage, even if fewer than four segments were successfully amplified and sequenced. Within-host variants on the OPV2 genome were identified with the R package deepSNV (64), using the OPV2 plasmid control to account for sequencing errors and strand bias. Minor iSNV (< 50% frequency) in the cohort samples were filtered using the following criteria: deepSNV p-value < 0.01, average mapping quality > 20, average Phred score > 35, and average read position in the middle 75% of the read (positions 31 and 219 for 250 bp pair reads). For samples sequenced in duplicate, we only used variants identified in both samples; we assigned frequency using the sample that had higher coverage at the site. We only identified iSNV present at a frequency of > 5%. Sites that were monomorphic after applying these filter criteria were assigned a frequency of 100%. The analytic pipeline was used to determine the position of each base in the coding sequence of the viral polyprotein and assign it as synonymous or non-synonymous relative to the sample consensus.

To obtain haplotype information specifically for VP1-143, codon frequency was identified by finding all reads that spanned the codon, filtering by MapQ > 20 and Phred score of each base > 20, counting the number of reads within each codon, and dividing by the number of reads that passed the quality filters. Samples with quality read depth less than 150 were excluded.

### Permutation test for parallel mutations

We quantified the probability that mutations would arise in parallel across a given number of individuals by implementing a permutation test. We first assumed that all sites were equally likely to mutate. We found the number of mutations that occurred in 83 individuals with variant-quality samples relative to the OPV2 reference above a frequency of 5%. We used this distribution to draw sites randomly across the genome, accounting for the length of the region amplified in our assay and excluding primer binding sites. We then found the number of sites shared by a given number of individuals. We ran this permutation 1000 times and calculated the p-value as the number of permutations with a number of shared sites equal to or greater than the observed data for a given group (e.g. mutations shared by two individuals, etc.). We simulated constraint on the mutability of genomic sites by restricting the fraction of sites available to mutate. We chose a fraction available of 60% to reflect the known distribution of fitness effects in poliovirus based on experimental data (65).

### Estimation of the transmission bottleneck

Models for estimating the transmission bottleneck were implemented as described in McCrone et al. 2018 (32). In the presence-absence model, we assessed whether donor iSNV are found in the recipient. We assumed perfect detection of transmitted iSNV and that the probability of transmission of donor iSNV are determined by the measured frequency at the time of sampling. We modeled the probability of transmission as a binomial sampling process, depending on the donor iSNV frequency and the bottleneck size (N_b_). We used maximum likelihood optimization to estimate the bottleneck size distribution, assuming bottlenecks across pairs follow a zero-truncated Poisson distribution. In the beta-binomial model, we relaxed the assumption of perfect detection of iSNV in the recipient by accounting for false-negative variant calls and stochastic loss below our detection threshold. We use our benchmarking data to supply the sensitivity of variant identification by frequency and titer, rounding down to the nearest titer threshold (e.g. 4.5 × 10^4^ copies/μL, 9 × 10^3^ copies/μL, etc.). We assume sensitivity in each range is the same as that of the titer threshold.

### Data availability and analysis code

The raw sequence reads for the Matlab samples and the benchmarking experiment are available on the NCBI Sequence Read Archive in BioProject PRJNA637613. Reads aligning to the human genome were filtered out by the SRA. The code for the primary analysis of within-host variants is publicly available at https://github.com/lauringlab/variant_pipeline. The rest of the code for data analysis and generation of the figures was written in R version 3.5.0 and python2.7 and is publicly available on GitHub at https://github.com/andrewvalesano/Poliovirus_Intrahost.

## Acknowledgements

We acknowledge and thank the participants and field workers in the original vaccine trial for their many contributions. We acknowledge the following individuals for their roles in implementation of the vaccine trial and sharing of samples: Dr. Mohammed Yunus, M.B.B.S. (Division of Infectious Diseases, icddr,b, Dhaka, Bangladesh), and Dr. William A. Petri, M.D./Ph.D (Division of Infectious Diseases and International Health, University of Virginia, Charlottesville, VA). We thank Suzanne Stroup for technical support. We thank Andrew Macadam, PhD and Cara Burns, PhD for generously providing the OPV1 and OPV2 plasmids, respectively. We thank JT McCrone for helpful discussion on data analysis. This work was funded by a grant from the Bill and Melinda Gates Foundation (to MF and MT) and a Burroughs Wellcome Fund Investigator in the Pathogenesis of Infectious Diseases Award (to ASL).

---

**Supplementary Table 1:**
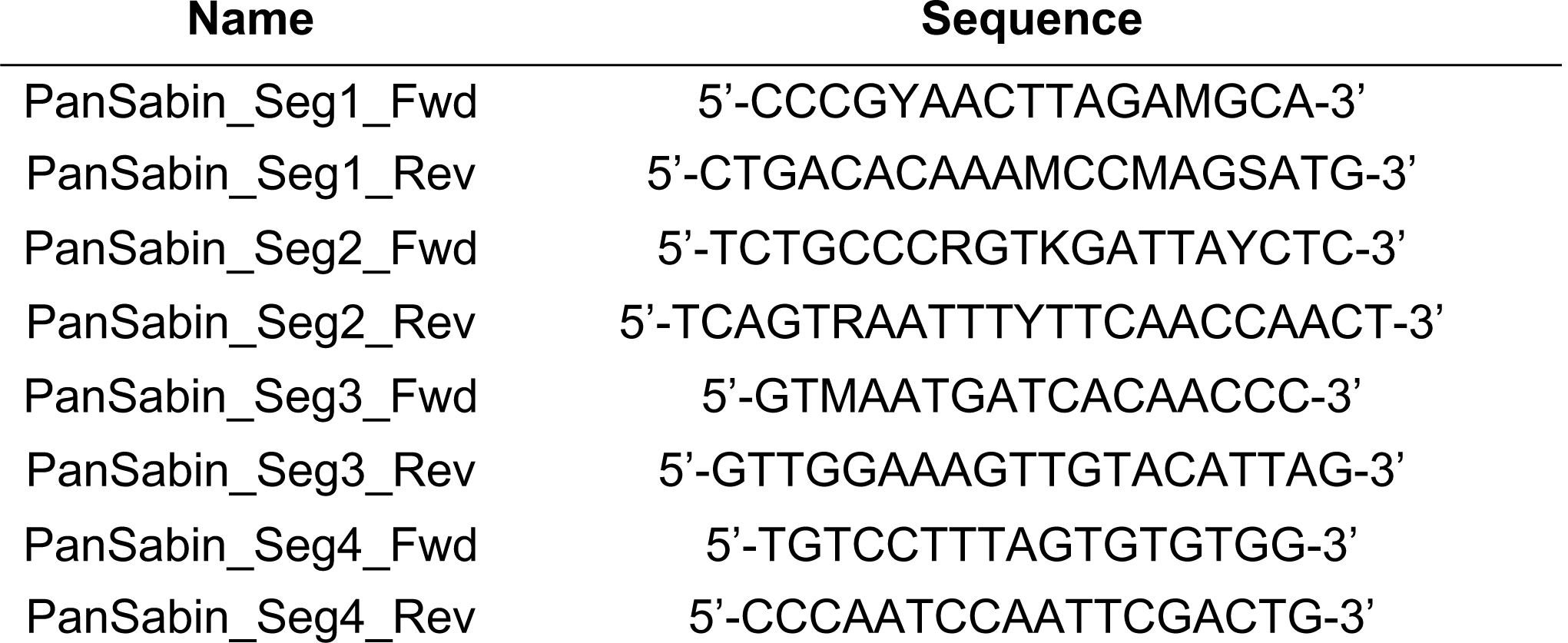
Genome amplification primers used in this study.

**Supplementary Table 2:** Gene-wise estimates of dN/dS ratio.

| Gene | Omega (dN/dS) |
| --- | --- |
| VP4 | 0.26326 |
| VP2 | 0.05951 |
| VP3 | 0.44309 |
| VP1 | 1.20709 |
| 2A | 0.25478 |
| 2B | 0.00010 |
| 2C | 0.05137 |
| 3A | 0.09244 |
| 3B | 0.00010 |
| 3C | 0.10622 |
| 3D | 0.04747 |

**Supplementary Table 3:** Samples from transmission pairs used in bottleneck analysis.

| **Donor ID<sup>a</sup>** | **Recipient ID<sup>a</sup>** | **Donor**<br>**Vaccination**<br>**Date<sup>b</sup>** | **Donor**<br>**Sample Date<sup>c</sup>** | **Recipient**<br>**Sample Date<sup>d</sup>** | **Time**<br>**Difference**<br>**(days)** |
| --- | --- | --- | --- | --- | --- |
| 115 | 10115 | 2016-01-26 | 2016-02-02 | 2016-02-01 | 1 |
| 171 | 10171 | 2016-01-25 | 2016-01-31 | 2016-01-31 | 0 |
| 702 | 20702 | 2016-01-25 | 2016-02-08 | 2016-02-08 | 0 |
| 927 | 10927 | 2016-01-28 | 2016-02-24 | 2016-02-17 | 7 |

<sup>a</sup> Anonymous IDs per individual.
<sup>b</sup> Date of mOPV2 administration in trial vaccination campaign.
<sup>c</sup> Date of sample collection from mOPV2 recipient used in the bottleneck analysis.
<sup>d</sup> Date of sample collection from the household contact used in the bottleneck analysis. For each recipient, this is the first longitudinal sample positive for OPV2 by RT-PCR.

**Supplementary Table 4:** Transmission bottleneck estimates for two models.

| Pair ID | Presence-absence model estimate<sup>a</sup> | Beta-binomial model estimate<sup>a</sup> |
| --- | --- | --- |
| 115 | 1 (1 – 2) | 1 (1 – 3) |
| 171 | 2 (2 – 4) | 2 (2 – 7) |
| 702 | 2 (2 – 2) | 2 (2 – 3) |
| 927 | 2 (2 – 5) | 2 (2 – 4) |

<sup>a</sup> 95% confidence interval shown in parentheses.

## Supplementary Figure Legends

**Supplementary Figure 1.**
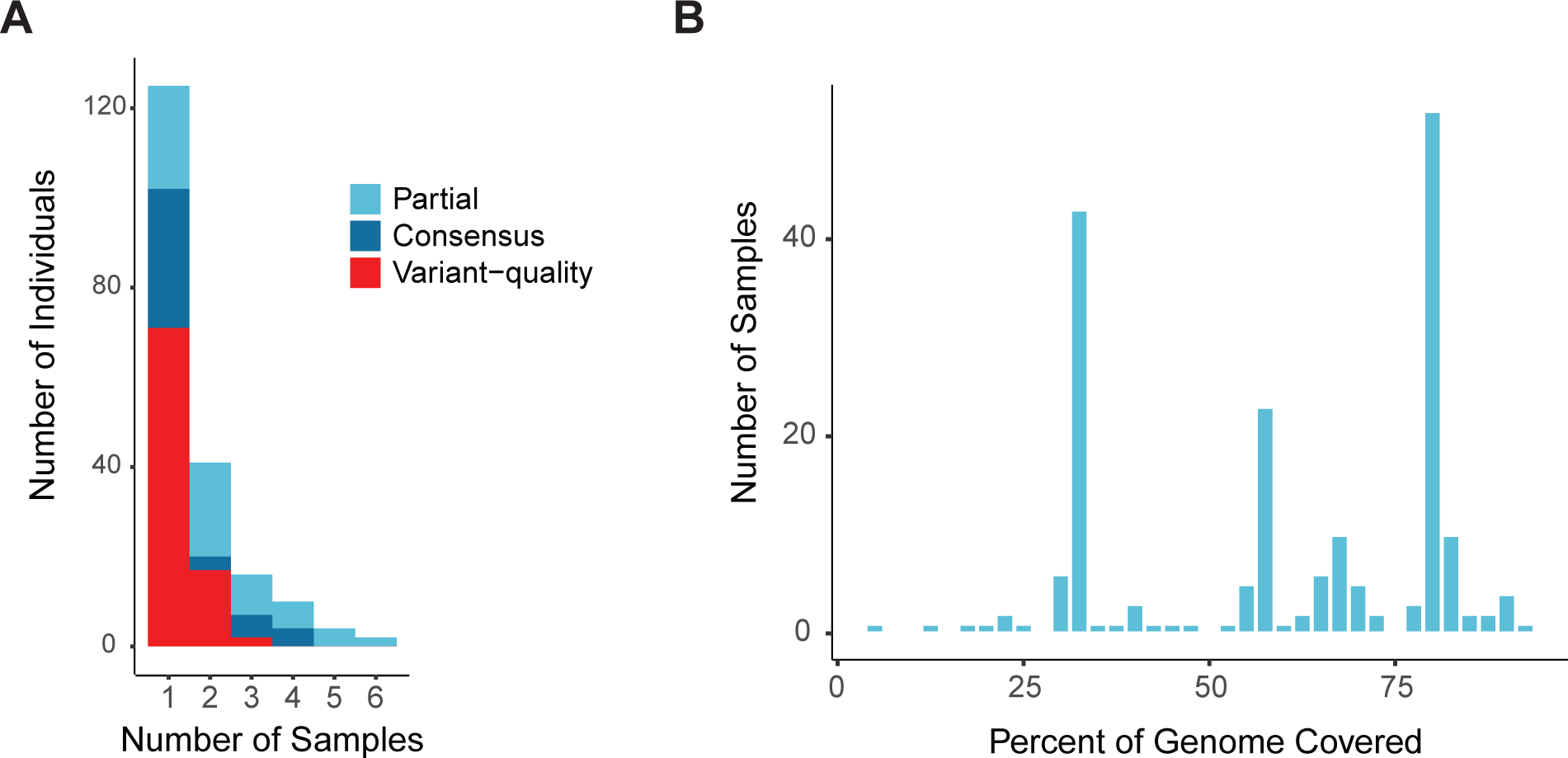
(A) Overlapping bar chart of the number of individuals (y-axis) by the number of samples sequenced from a given individual (x-axis). Colors represent the genome coverage groups shown in Figure 1. (B) Composition of the partial genome samples. Number of samples (y-axis) by the percent of the genome covered above a 10x threshold (x-axis).

**Supplementary Figure 2.**
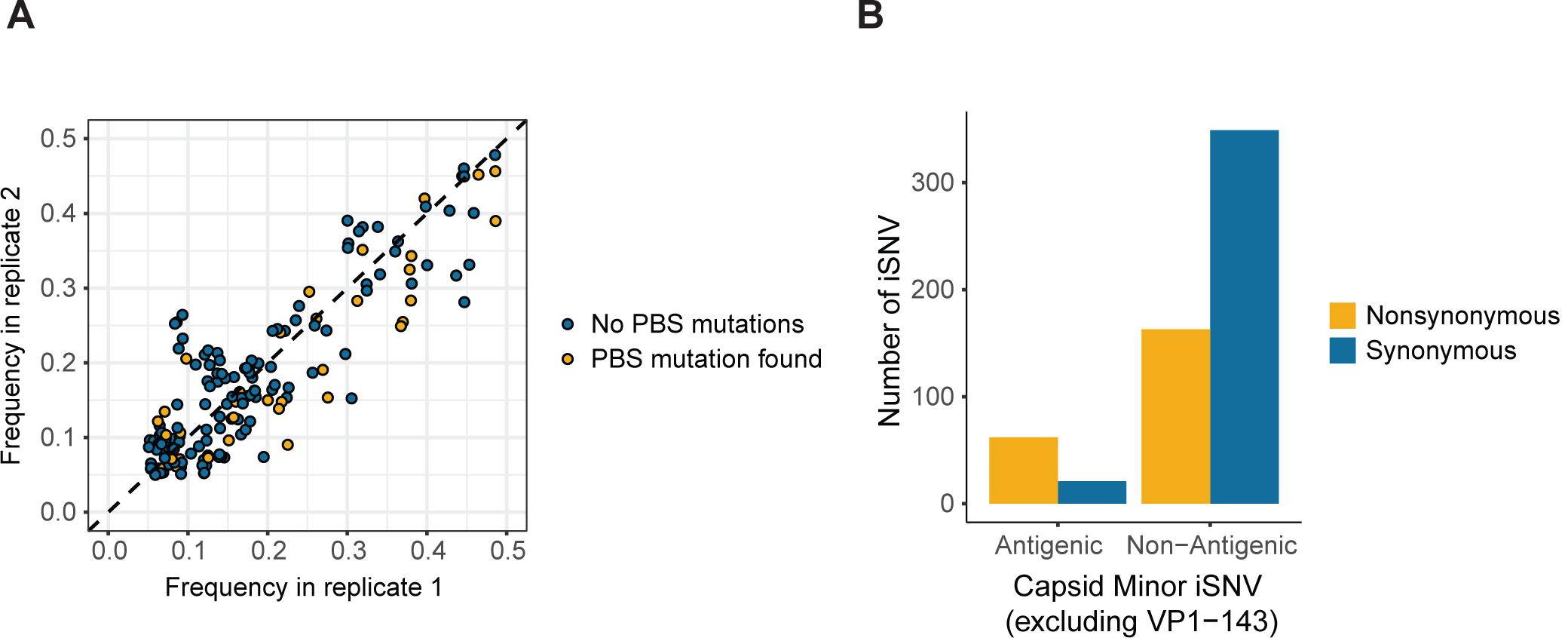
(A) Concordance of iSNV frequency measurements across 11 samples sequenced in duplicate. Frequency of iSNV in replicate 2 (y-axis) is shown by the frequency of an iSNV in replicate 1 (x-axis), with colors showing iSNV on amplicon(s) with or without mutations in primer binding sites. (B) Histogram of minor iSNV in the capsid region by antigenic status, excluding VP1-143. Nonsynonymous iSNV are shown in yellow, and synonymous iSNV in dark blue.

**Supplementary Figure 3.**
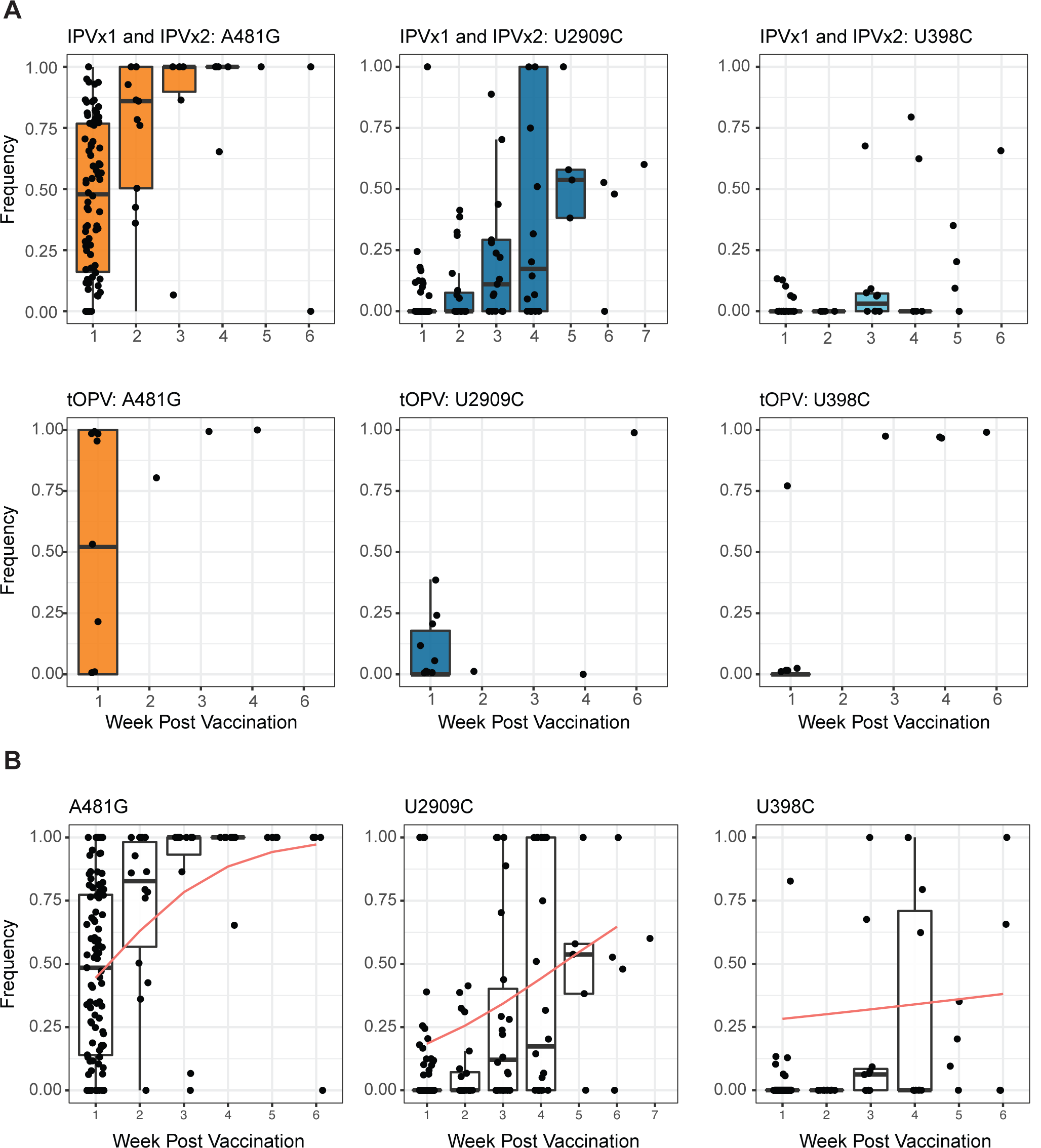
(A) Frequency of A481G, VP1-143X, and U398C by time from vaccination across arms of the vaccine trial. Samples from IPV arms are shown on the top, and samples from tOPV arms are shown on the bottom. Each point represents one sample, and boxplots are shown for weeks with five or more data points. Boxplots represent the median and 25^th^ and 75^th^ percentiles, with whiskers extending to the most extreme point within the range of the median ± 1.5 times the interquartile range. (B) Frequency of A481G, VP1-143X, and U398C by time from vaccination across with the beta regression model fits for each mutation (red lines). The underlying data are the same as in Figure 3A. Each point represents one sample, and boxplots are shown for weeks with five or more data points. Boxplots represent the median and 25^th^ and 75^th^ percentiles, with whiskers extending to the most extreme point within the range of the median ± 1.5 times the interquartile range.

**Supplementary Figure 4.**
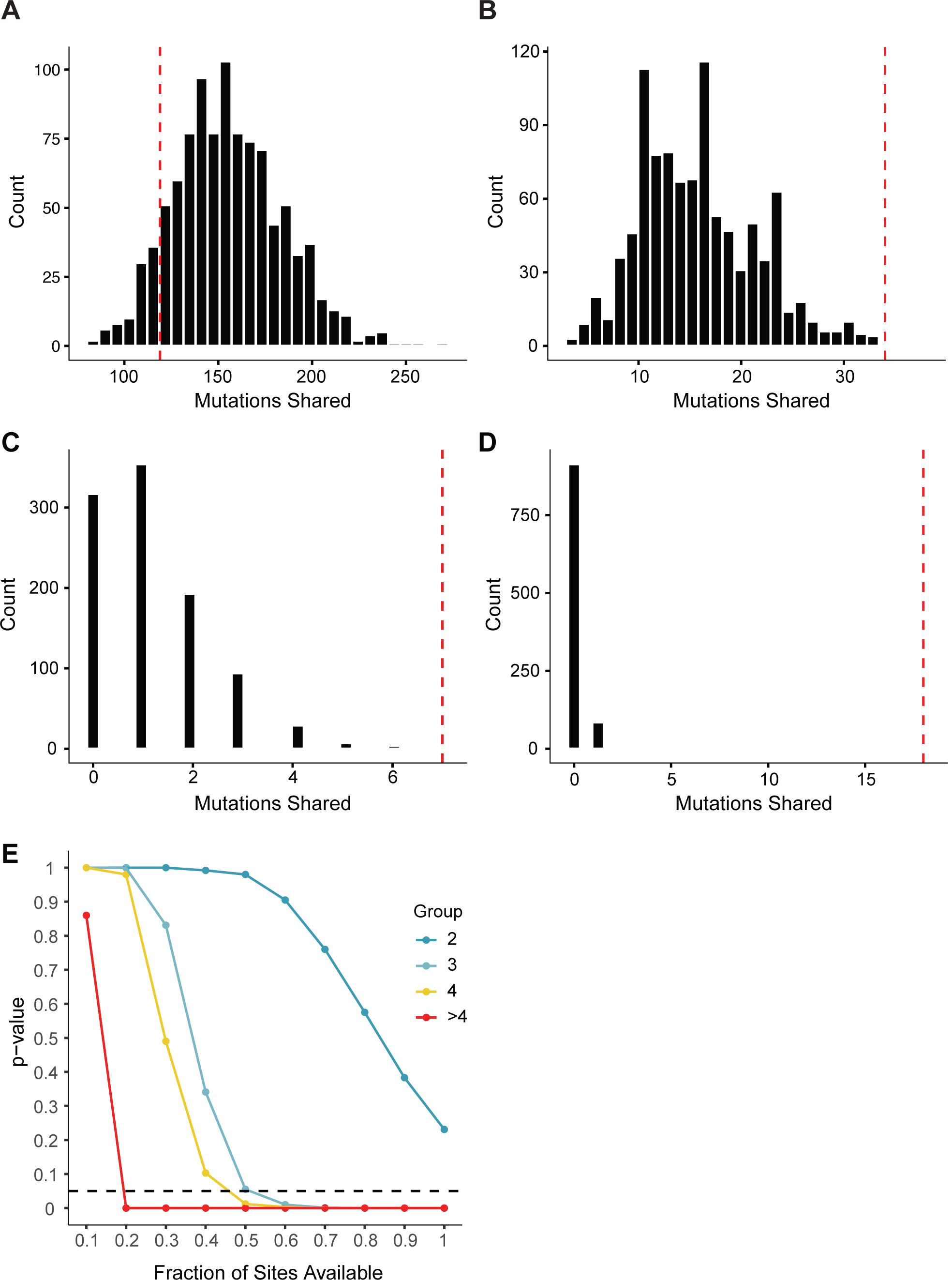
Results of permutation test for assessment of parallel mutations. In A - D, the dotted red line is the observed number of mutations shared across a given number of individuals, and the bars show the results of 1000 random permutations. Results are shown for mutations shared across two individuals (A), three individuals (B), four individuals (C), and greater than four individuals (D). P-values are designated as the proportion of random permutations equal or greater than the observed number of shared mutations. In E, the p-values for each group are shown as a function of the genome fraction available for mutations. The horizontal dotted line represents α = 0.05.

**Supplementary Figure 5.**
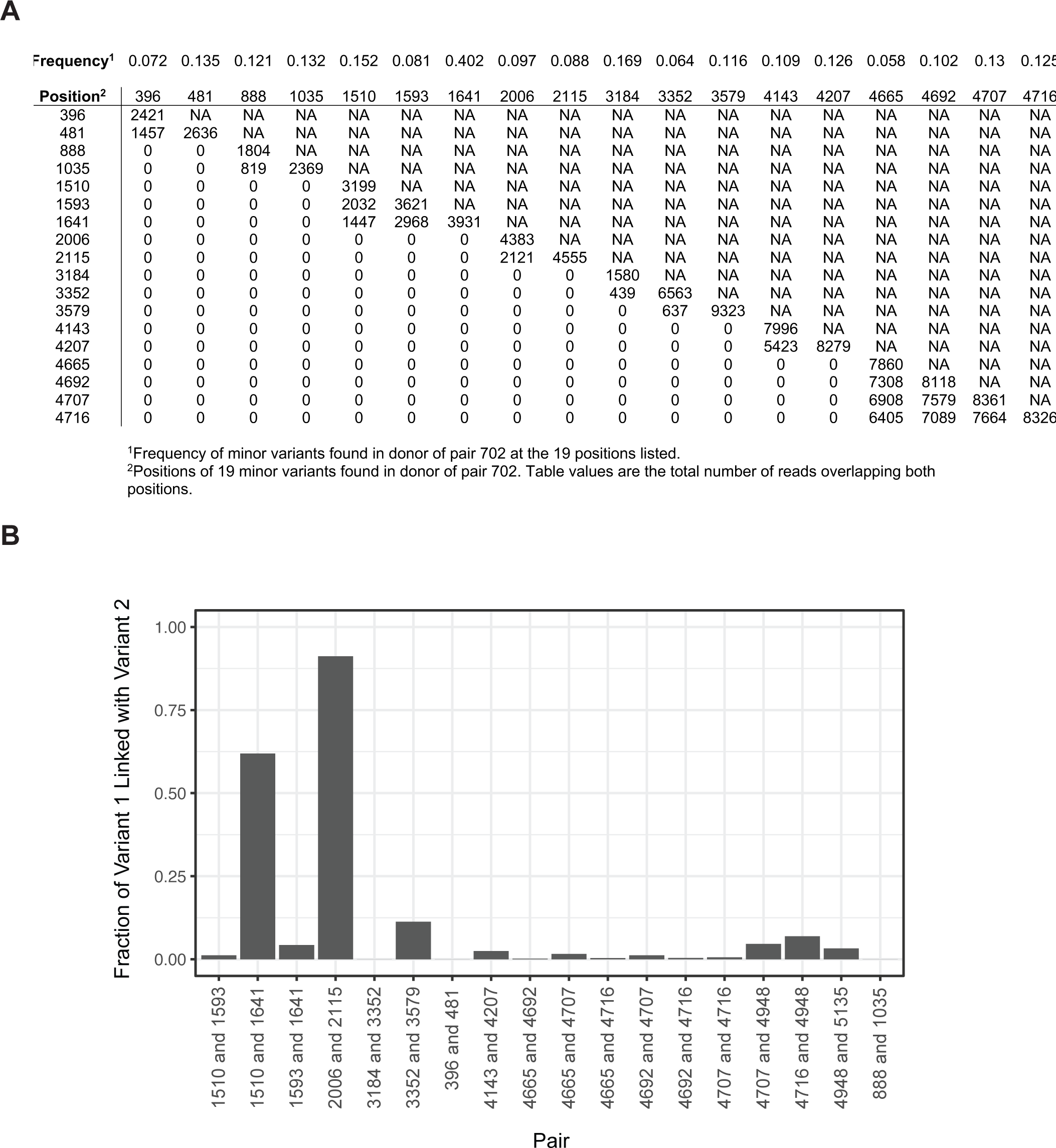
(A) The frequency of 20 minor variants present in the donor for pair 702 (top). The table values show the number of sequence reads overlapping each pair of minor variants. (B) Bar chart showing the fraction of minor variant 1 found linked to minor variant 2 in overlapping sequence reads. The 18 pairs of minor variants are shown here by their genome position.

